# Systematic identification of cooperation between DNA binding proteins in 3D space

**DOI:** 10.1101/036145

**Authors:** Kai Zhang, Nan Li, Wei Wang

## Abstract

Cooperation between DNA-binding proteins (DBPs) such as transcription factors and chromatin remodeling enzymes plays a pivotal role in regulating gene expression and other biological processes. Such cooperation is often via interaction between DBPs that bind to loci located distal in the linear genome but close in the 3D space, referred as trans-cooperation. Due to the lack of 3D chromosomal structure, identification of DBP cooperation has been limited to those binding to neighbor regions in the linear genome, referred as cis-cooperation. Here we present the first study that integrates protein ChIP-seq and Hi-C data to systematically identify both cis- and trans-cooperation between DBPs. We developed a new network model that allows identification of cooperation between multiple DBPs and reveals cell type specific or independent regulations. Particularly interesting, we have retrieved many known and previously unknown trans-cooperation between DBPs in the chromosomal loops that may be a key factor for influencing 3D chromosomal structure. The software is available at http://wanglab.ucsd.edu/star/DBPnet/index.html.

## Introduction

DNA binding proteins (DBPs) such as transcription factors (TFs), insulators and chromatin remodeling enzymes play key roles in many important biological processes. These proteins rarely function alone but rather cooperate with one another to regulate gene expression, epigenetic modifications and formation of 3D interactions between distal genomic loci[1, 2]. Identification of DBP cooperation is thus critical for understanding the mechanisms regulating these crucial molecular and cellular functions.

Previous studies have focused on identifying DBPs binding to neighbor genomic regions[3–7], which is hereinafter referred as cis-cooperation. Despite the great insight of these studies provided into revealing combinatorial regulation of DBPs, they missed cooperation between DBPs binding to distal genomic loci but localized in spatial proximity to form so called trans-cooperation. Trans-cooperation of DBPs either enhances the existing 3D contacts or creates new ones to bring functional elements such as enhancers to their target loci such as promoters. However, no study has thoroughly investigated the trans-cooperation and its relationship with cis-cooperation.

The ENCODE project has generated hundreds of ChIP-seq data to map binding sites of DBPs in multiple cell lines[8–10]. Recently, kilobase-resolution Hi-C data were available in two of these cell lines, GM12878 and K562[11]. These data provide an unprecedented opportunity to systematically map both cis- and trans-cooperation between DBPs. However, it is a great challenge to analyze this large amount of data and extract the cooperation among multiple rather than a pair of DBPs.

To tackle this challenge and comprehensively catalog DBP cooperation, we present here a new model to constructing networks that represent both cis- and trans-association between DBPs. Analyzing these networks in GM12878 and K562 has revealed complex cooperative relationship among TFs, histone modifications, chromatin remodeling enzymes and chromatin structure mediators. Through identification of modules and cliques in the network composed of trans-cooperation, we have uncovered many DBP interactions in the chromatin loop regions. Particularly interesting, many of these trans-cooperated DBPs directly interact with one another, which suggests their binding may be important for loop formation or stabilization in 3D space. Comparative analysis between GM12878 and K562 revealed cell-type specific cooperation between DBPs that are critical for regulating cell-type specific functions.

## Results

### Gaussian Graphical model (GGM)

To systematically identify DBP cooperation, we analyzed DBP ChIP-seq data using Gaussian graphical model (GGM)[12]. GGM is an undirected probabilistic graphical model with the assumption that the data follows a multivariate Gaussian distribution with mean μ and covariance matrix Σ. Let Σ^−1^ be the inverse of covariance matrix. If the ij*th* component of Σ^−1^ is zero, then variable i and variable j are conditional independent given all other variables in the network[12]. This important theorem serves as the foundation for GGM to infer direct interactions from data. Unlike relevance networks or correlation networks[13–16], in which edges are determined based on marginal correlations, GGM provides stronger criterion of dependency, and thus further reduces false positive rate[17]. For this reason, GGM has been applied to constructing biological networks[18, 19]. However, classic GGM includes too many edges in the estimated graph, which also raises an issue of overfitting the data. To cope with this, Friedman et al. proposed an efficient algorithm, named graphical Lasso, to solve this problem[20]. Recently, Liu et al. developed a data transformation method called Copula that can be used with the graphical Lasso algorithm to relax the assumption of normality in constructing GGM[21, 22]. Based on these recent advancements, we developed a new method to systematically investigate cooperation between hundreds of DBPs.

Before applying to the DBP ChIP-seq data, we assessed the performance of GGM using synthetic data. Firstly, we generated an Erdős-Rényi random graph (see Methods). To generate samples according to the simulated graph, we built a covariance matrix of the multivariate Gaussian distribution by assigning each ijth component a non-zero covariance if node i and j are connected in the random graph. All other components are set to zero. We then drew samples from a multivariate Gaussian distribution with zero mean and the constructed covariance matrix. These samples were then used as input to the network construction algorithms. For comparison, we selected ARACNE[16], a popular algorithm for constructing gene regulatory networks. It uses an information theory approach to eliminate most indirect interactions in networks inferred by co-expression methods, and has been proved useful by independent studies[23–27]. We generated 10 networks with 50 nodes and another 10 networks with 100 nodes. We observed superior performance of GGM with an average AUC 0.923, obviously higher than that of ARACNE (AUC=0.822) (Fig. 1a). This simulation showed that, when experimental data follows Gaussian distribution, our GGM can precisely reconstruct the underlying graphical model. Because the DBP ChIP-seq data can be noisy and may not be Gaussian distributed, we adopted the Copula algorithm[21, 22] for data transformation to approximate the real distribution. To assess the GGM performance in this scenario, we generated synthetic gene expression profiles using GeneNetWeaver 3.1[28], an in silico simulator that employs dynamic model to simulate gene regulatory networks. The ground truth were subnetworks taken from yeast gene regulatory network with size 50 and 100, respectively. For each size, we performed 10 different simulations (network files and sample data are provided in the Supplementary Data). Again, GGM outperformed ARACNE (Average AUC 0.695 vs 0.615) (Fig. 1b).

**Figure 1.**
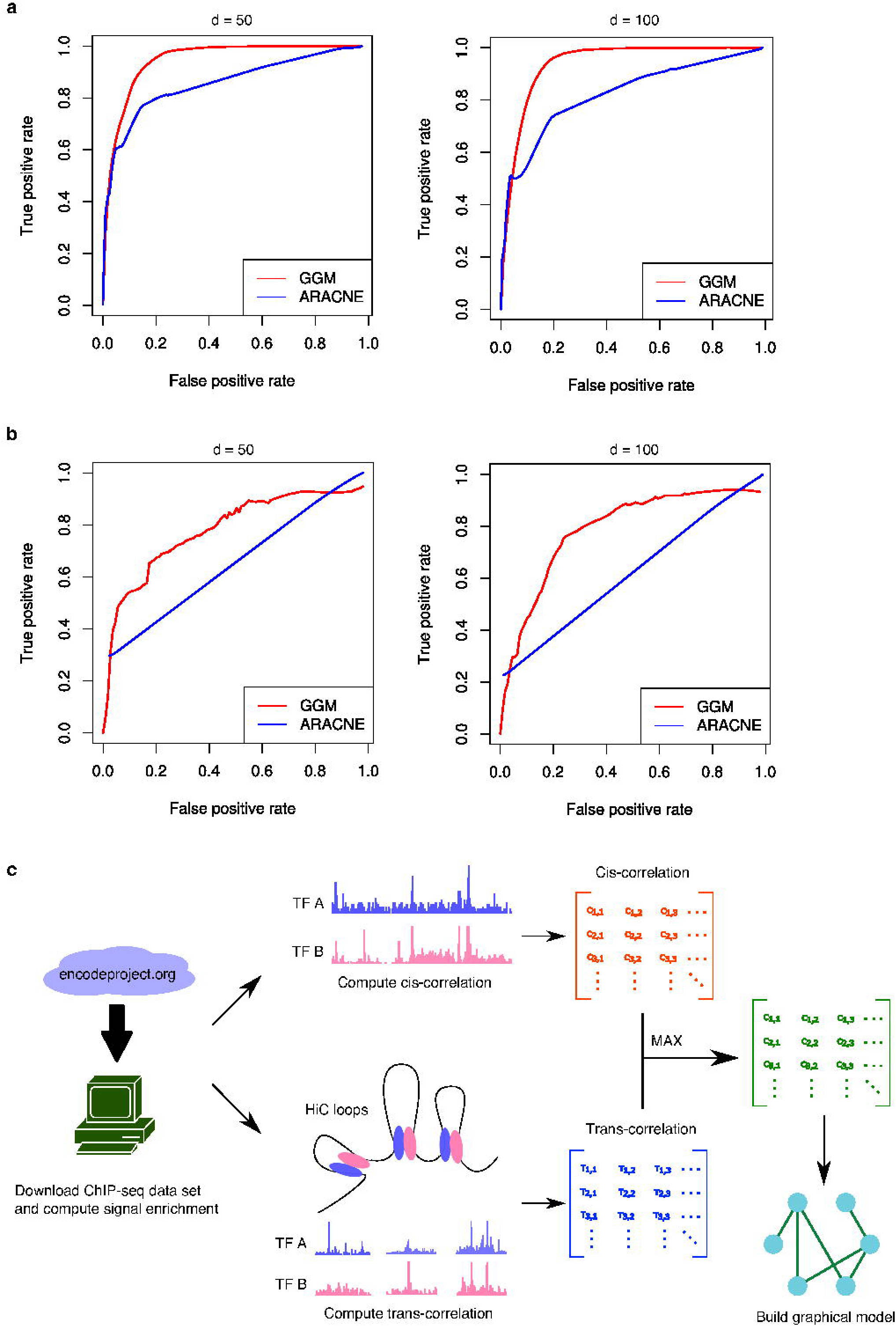
The performance of GGM is consistently better than ARACNE. Each plot shows the average curve from 10 independent simulations. (a) ROC curve for samples generated from random networks. For each simulation 500 (Left) or 1000 (Right) samples were generated from a network of 50 (Left) or 100 (Right) nodes. (b) ROC curve for samples generated from yeast sub-networks. For each simulation 500 (Left) or 1000 (Right) samples were generated from a network of 50 (Left) or 100 (Right) nodes. (c) Workflow of the DBPnet pipeline.

It is worth of noting that GGM is much faster than ARACNE when the sample size is large. The time complexity for ARACNE is O(N^3^+N^2^M^2^), where N is the number of variables or nodes in the network, M is the number of samples; as it scales with M^2^, it is not suitable for our application where we have more than 10,000 samples (the number of ChIP-seq peaks). In contrast, GGM, with a time complexity O(N^3^+N^2^M), can easily handle a large number of samples. In practice, we observed 50 ~ 100 times faster of GGM than ARACNE on the synthetic data sets (Supplementary Table 1).

### Constructing DBP cooperation network

After we confirmed the GGM performance on constructing gene network from synthetic data, we applied it to DBP ChIP-seq and Hi-C data, aiming to systematically detect DBP cooperation (Fig. 1c). We considered both cis (DBPs bind in nearby linear genome) and trans (DBPs bind to loci that are spatially close but linearly distal in the genome) cooperation between DBPs. We first computed cis and trans correlation score for each pair of DBPs separately using the 84 DBP and six histone modification (H3K4me1, H3K4me3, H3K9me3, H3K27ac, H3K27me3, H3K36me3) ChIP-seq as well as loops called by the 5kbp-resolution Hi-C data in a lymphoblastoid cell line GM12878[29] (see details in Methods). We then merged cis and trans correlation matrices by keeping the larger correlation score at each entry. This matrix was used to construct GGM, which represents the DBP cooperation network.

The GGM network (Fig. 2a) identified 484 associations among the DBPs. An edge between two proteins may indicate either a direct physical interaction or a co-occurrence of binding sites without direct interaction. To examine whether our model can recover direct protein-protein interactions, for each edge we searched for supporting evidence from the public protein-protein interaction databases (Methods). Remarkably, 11% of edges (p-value is 10e-9) in the GGM network are also present in the PPI network (Fig. 2b). Another 80% of the associated DBPs are separated by one protein in the PPI network (the intermediate protein may not be analyzed by ChIP-seq). These evidences strongly supported that the DBP associations recovered by the GGM network are reliable and likely represent direct physical contacts.

**Figure 2.**
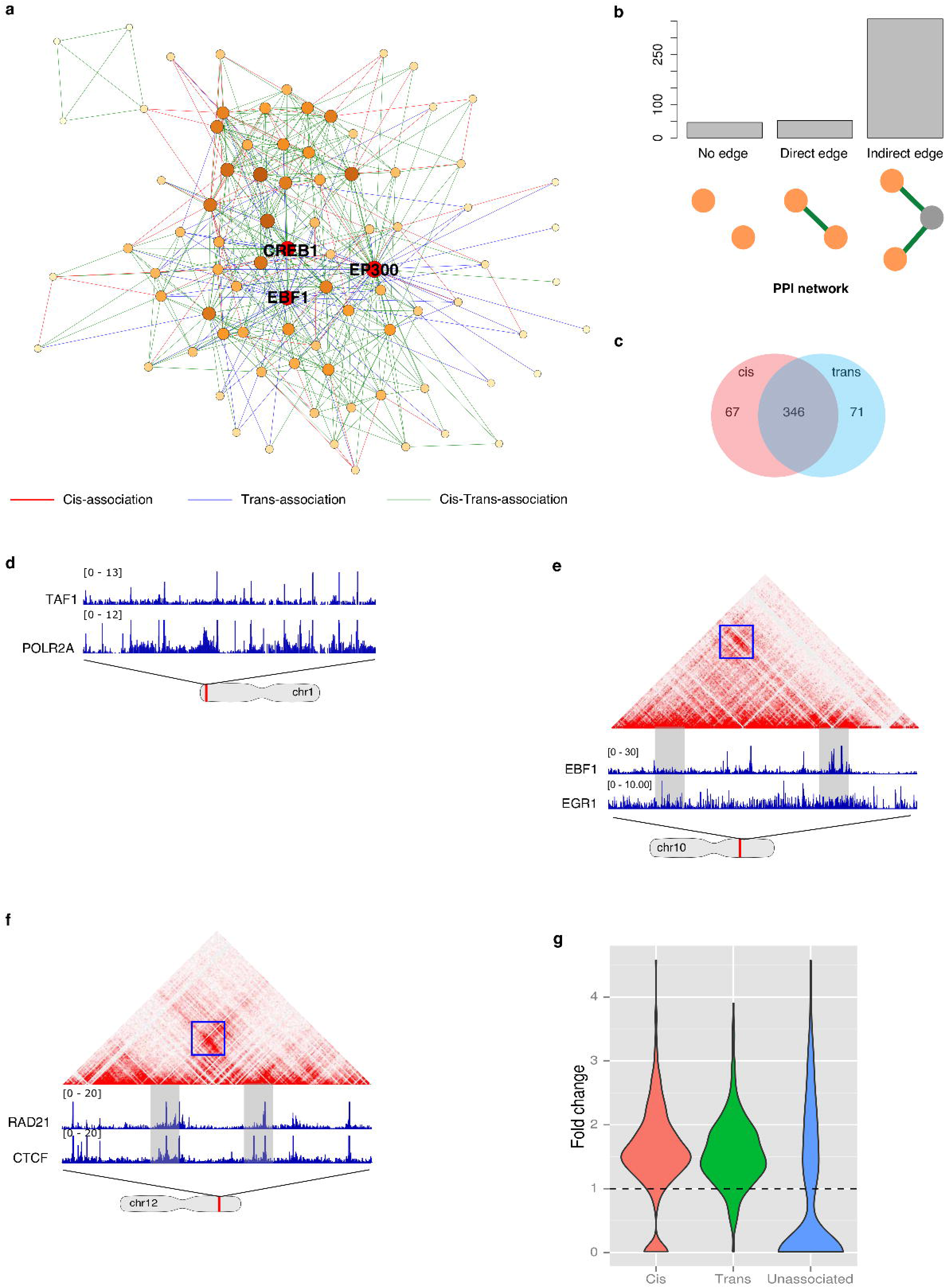
Constructing DBP cooperation network in GM12878. (a) DBP cooperation network in GM12878, with network hubs (EP300, EBF1, CREB1) being highlighted. (b) A significant portion of DBP cooperation is supported by the evidences of direct protein-protein interaction. (c) The majority of DBP cooperation is a mixture of cis and trans cooperation. (d) An example of cis-cooperation. (e) An example of trans-cooperation. (f) An example of mix cooperation. (g) Disease associated CVs are enriched in cis- and trans cooperative sites.

To characterize the topological properties of the DBP cooperation network, we first plotted the node degree distribution, which follows power law, indicating that it is a scale-free network[30] (Supplementary Fig. 1). A property of scale-free network is the existence of hubs, the highly connected nodes critical for network stability. To identify hubs, we ranked the nodes in our network by two popular metrics used to measure the importance of a given node in a network - node degree and eigenvector centrality. Node degree, the number of links a node has, is the simplest centrality metric. A more sophisticated metric is the eigenvector centrality which assigns high centralities to nodes that are linked to many other well-connected nodes[31]. We ranked the nodes by their node degree and eigenvector centrality. EP300, CREB1 and EBF1 as the top three DBPs that have the best average rank (Fig. 2a and Supplementary Table 2). EP300 is an important co-factor that cooperates with many TFs[32, 33] and CREB1 plays a central role in immune system through binding to c-AMP response element, a ubiquitous DNA motif, to regulate gene transcription[34, 35]. It is not unexpected that these two general DBPs were found as hubs. On the other hand, EBF1 (Early B-cell Factor 1) is mainly expressed in B-lymphocyte (GM12878 is a lympoblastoid cell line) and is pivotal for maintenance of B cell identity[36]. Moreover, DBPs that are highly correlated with EBF1 include EP300, PAX5, SP1, BHLHE40, TCF12, and BCL11A. EP300 and SP1 are general transcriptional activators. The functions of PAX5, TCF12 and BCL11A are highly specific to B-lymphocyte development[37–40].

### Identifying cis and trans interactions between DBPs

DBPs can cooperate through cis or trans interactions, which can be determined for each DBP pair using the constructed network. In this study we defined an interaction as cis or trans cooperation if the cis or trans correlation score is larger than a pre-selected cutoff (0.3, see Methods). In GM12878, we found roughly same numbers of cis and trans edges, 413 and 417 respectively. We noticed a great overlap between cis and trans edges (Fig. 2c). DBP cooperation falls into three categories: cis dominant, trans dominant and mix (Supplementary Table 3).

Cis dominant association between two DBPs represents a strong occurrence in linear space but not in long-range interacting loci that form loops in the 3D space. In this category we recovered some previously known interactions such as RNA Pol II-TAF1 interaction[41] (Fig. 2d). Interestingly, we found 71 trans-dominant edges in GM12878. Most of these edges show weak cis correlation but have significantly larger trans correlations such as EP300-MYC whose cis and trans correlation scores are 0.127 and 0. 348 (z-score: −0.081 and 1.301). Indeed, a number of independent studies have shown that EP300 and MYC can cooperate to regulate gene transcription and the physical interaction between them was previously reported[42–44]. We also identified novel DBP cooperation mediated by trans-interactions. For example, EBF1 is an important TF in B-lymphocyte, and Egr-1 is one of the important transcriptional regulators induced upon B cell antigen receptor activation[45]. Both EBF1 and Egr-1 have crucial roles in B cell development and differentiation. However, the interplay between these two proteins has not yet been reported. In our network, we found a trans-dominant edge between EBF1 and Egr-1 (Fig. 2e), which suggests that they may form long-range loops to regulate cell type specific genes (4.4% loop regions contain peaks of both EBF1 and Egr-1). Therefore, our framework provides a systematic way to uncover trans interactions between DBPs that are otherwise impossible to be identified using previous approaches. The mix category consists of DBP pairs with both cis and trans associations. Most associations fall into this category. A well-known example is CTCF-RAD21 (Fig. 2f). Previous studies have focused on identification of cis associations between DBPs[4]. However, the last two categories of DBP associations can only be identified through the integration of DBP ChIP-seq and Hi-C data.

To further confirm the importance of DBP cooperation, we carried out genotype variations (GVs) enrichment analysis in regions bound by cooperative DBPs. Firstly, we downloaded disease associated GVs from the NHGRI-EBI GWAS database[46]. Assuming A and B are two cis-cooperative DBPs, we then calculated enrichments of GVs at two types of binding sites: (1) sites bound by both A and B; (2) sites bound by A or B. The ratio of the former to the latter is termed as the fold change of GV enrichment for cis interacting sites. Likewise, to calculate this fold change for trans interacting sites, we required A and B are trans-associated. The GVs enrichment in sites involved in trans association are then computed and the fold change is calculated similarly. Fig. 2g shows that the vast majority of DBP associations, either cis or trans, are more enriched with disease associated GVs (p-values are 1.5e-21 and 8.6e-27 respectively compared to unassociated DBP pairs), which suggests that DBP cooperation has important functional implication in a variety of diseases. Therefore, we anticipate that the binding sites of cooperative DBPs can be used to prioritize GVs to identify causal associations.

### Identifying DBP modules

To identify cooperation between multiple DBPs, we searched for modules in the constructed network. Previous approaches have focused on cis DBP modules but trans DBP modules have not been systematically identified. The DBP cooperation network provided an unprecedented opportunity to tackle this challenge. We first used community detection algorithm[47] to identify 4 communities in GM12878 (Fig. 3a and Supplementary Table 4). A community is composed of tightly interconnected DBPs[47]. DBPs within a community presumably have related functions. For instance, we found that the yellow community consists of CTCF, RAD21, ZNF143, SMC3 and YY1. Because these proteins are important to mediate chromatin looping[29, 48–50], the main function of the yellow community is likely to be maintenance of chromatin structure.

**Figure 3.**
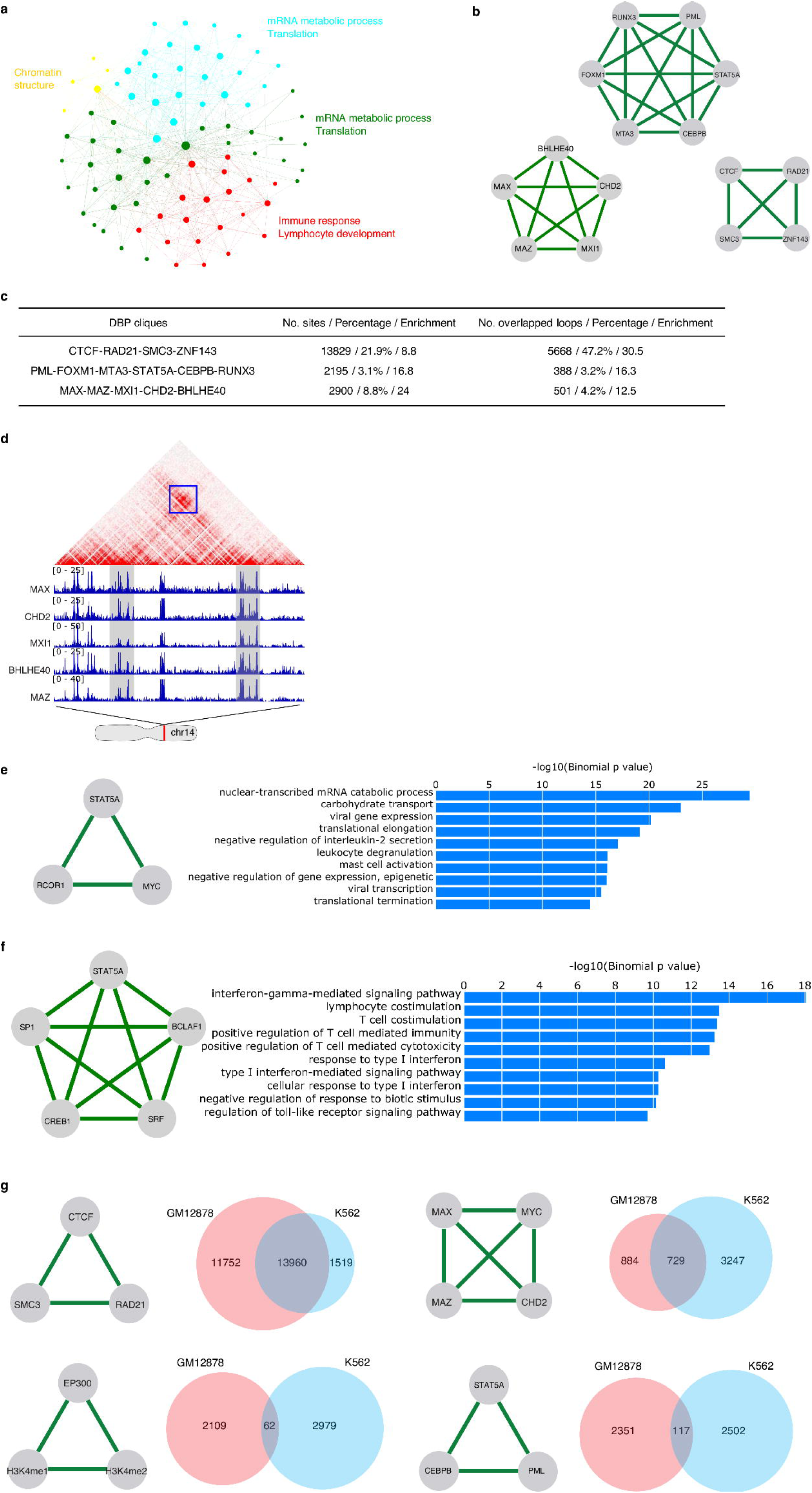
Network analysis reveals functions of DBP modules in GM12878 and K562. (a) Communities in DBP cooperation network and their functions. (b) Top DBP cliques. (c) The member DBPs in a clique frequently co-occur in a cis and trans fashion. The central column gives the number of regions bound by all DBP members in a clique, the percent of these regions and its fold enrichment over background. The right column gives the number of loops that are overlapped with the DBP binding sites, its percent and fold enrichment over background. (d) An example of DBP cliques. (e) An example of K562 specific DBP cliques and enriched GO terms of its binding sites. (f) An example of GM12878 specific DBP cliques and enriched GO terms of its binding sites. (g) Top conserved DBP modules in K562 and GM12878.

Interestingly, all the six histone modifications are located in the green community, including H3K4me1/2/3, H3K9ac, H3K27ac, H3K36me3, which indicates the global correlation between histone modifications despite their distinct roles in marking regulatory elements. To reveal the functions of different communities, we analyzed the ChIP-seq peaks of proteins in this community. We ranked genomic loci by the number of proteins bind to them and selected the top 5000 loci as input to GREAT analysis searching for enriched GO terms. We found the green community is linked to mRNA metabolic process and translation related functions (Supplementary Fig. 2a).

Similarly, the cyan community is enriched with general GO terms such as “mRNA metabolic process” and “translation regulation” (Supplementary Fig. 2b), which suggests both green and cyan community are involved in regulating the basic functionality of the cell. In fact, these two communities contain numerous general transcription machinery proteins such as RNA polymerase II, TATA box binding protein (TBP) and TAF1.

In contrast to the general function of the green and cyan communities, the most enriched GO terms for the red community are “immune response”, “leukocyte activation”, and “lymphocyte activation” that are highly related to B cell functions (Supplementary Fig. 2c), which suggests that this community plays important roles in determining cell type specificity. Indeed, many proteins in this community are known to be important for immune system development, such as STAT5A[51], BATF[52], BCL3[53], and RELA[54].

### Identifying DBP cliques that potentially form protein complexes

DBPs often cooperate with one another to form a large complex. Identification of such complex is crucial for understanding the mechanisms of transcriptional regulation. Therefore, we searched for maximal cliques in the network. In a clique, each node is linked to all the other nodes. DBPs involved in a clique are thus likely to form a protein complex. In GM12878, we identified 220 cliques (Supplementary Table 5). We ranked DBP cliques by their average correlation scores of each DBP pair. We observed that most of the top cliques have both high cis and trans interaction edges, which suggests that they are likely to form complexes mediating chromosome loop formation. Fig. 3b shows the top 3 highest ranked cliques. Next, we checked the percentage of shared peaks in the union of all DBP peaks and found the loops in these shared peaks. All the DBPs in the cliques share a significant amount of peaks that occur in loops (Fig. 3c), which confirmed the co-occurring bindings of the DBPs in a clique.

One of the top ranked cliques is CTCF-RAD21-SMC3-ZNF143. RAD21 and SMC3 are components of cohesin complex. Cohesin is a multi-subunit protein complex and plays an essential role in sister chromatid cohesion and chromosome segregation during cell division[55]. Cohesin is also crucial for regulating gene expression and mediating chromatin long-range interaction[56]. Cohesin-dependent chromatin interactions are usually mediated by the cooperation of cohesin and CTCF[49, 50]. The involvement of Zinc Finger Protein 143 (ZNF143) in this complex has also been reported[48, 49]. ZNF143 is believed to provide sequence specificity for chromatin interactions[57]. Overall, our analysis successfully recovered this important and well characterized loop forming complex.

The other two cliques, PML-FOXM1-MTA3-STAT5A-CEBPB-RUNX3 and MAX-MAZ-MXI1-CHD2-BHLHE40 (Fig. 3d), are novel and their functions have not been characterized. STAT5A and RUNX3 are two of the major transcription factors that play essential roles in lymphocyte development. The physical interaction between STAT5 and RUNX3 has been reported[58]. Moreover, CEBPB binds to RUNX2 which has been shown to be associated with RUNX3[59, 60]. To investigate the function of this module, we extracted all loci bound by these six DBPs and performed GREAT analysis. The most significant GO terms are “immune response”, “leukocyte activation”, and “lymphocyte activation”. These results suggested this novel module may play important roles in the immune system development of lymphocytes.

The functions of MAX-MAZ-MXI1-CHD2-BHLHE40 clique seem to be more general. GREAT analysis of their binding sites revealed enriched GO terms as “ribonucleoprotein complex biogenesis”, “nuclear-transcribed mRNA catabolic process ribosome biogenesis” and “translation”. We then searched for evidences supporting their cooperation in literature. The interaction between MAX and MXI1 is well studied[61] but interactions between other proteins have not been reported. However, the functions of these proteins are related. For example, BHLHE40 is a repressor that can interact with and recruit HDACs, which suggests a role of BHLHE40 in chromatin remodeling. CHD2 is also a chromatin remodeler. These observations suggest that DBPs in this clique may act together to alter chromatin states and regulate gene translation.

### Comparative analysis of DBP cooperation networks across cell types

DBPs have different cooperative modes in different cells. To perform a comparative analysis of the networks in different cell types, we focused on 68 proteins for which ChIP-seq data were available in both K562 and GM12878 and constructed TF cooperation networks in these two cell types.

In order to find cell type specific DBP cliques, we first identified cell type specific edges in the 68-node networks. We then searched for cliques in both GM12878 and K562 specific networks that consist of edges present in one but not the other cell line. We found 74 and 7 cell type specific cliques for GM12878 and K562, respectively (Supplementary Table 6). Cell type specific cliques shed light on how cells achieve transcriptional specificity through combinatorial regulation of DBPs. For example, STAT5A is a member of STAT protein family. It is activated by a number of cytokines and plays a central role in the development of many different organs. However, how STAT5A cooperates with other DBPs to carry out cell type specific regulation is largely unknown. Our analysis showed that STAT5A, together with MYC and RCOR1, forms a clique in K562, which is absent in GM12878. MYC is an oncogene and has been shown to play a critical role in leukemia formation[62, 63]. STAT5A-MYC cooperation may be important to maintain the state of leukemic cells. To further characterize the functions of STAT5A-MYC-RCOR1 module, we performed GREAT analysis on loci bound by all the three proteins in K562 and observed functions specific to leukocyte, such as “leukocyte degranulation”, “regulation of interleukin-2 secretion”, “mast cell activation” (Fig. 3e). These functions are drastically different from those enriched in GM12878 where STAT5A is associated with BCLAF1, SRF, CREB1 and SP1; GREAT analysis on the shared peaks suggests this module is involved in lymphocyte specific functions (Fig. 3f).

Next, we sought to identify common DBP modules in GM12878 and K562. Firstly, we extracted a common network using edges shared by the two networks. We then searched for cliques in this network. We identified CTCF-RAD21-SMC3, MAX-MYC-MAZ-CHD2, EP300-H3K4me1-H3K4me2 and STAT5A-CEBPB-PML as top ranked cliques (Fig. 3g). CTCF-RAD21-SMC3 interaction is known to be conserved across different cell types and it is not surprising that this clique is shared between the two cells. In the MAX-MYC-MAZ-CHD2 clique, MAX-MYC-MAZ is also a well-known complex that is found in multiple cell lines but their interaction with CHD2 has not been reported. The involvement of the chromatin remodeling gene CHD2 in MAX-MYC-MAZ complex suggests MAX-MYC-MAZ may utilize CHD2 to modify chromatin structure and alter gene expression. EP300-H3K4m1-H3K4me2 represents enhancer’s signature, and has been found in many cell types. In the clique of STAT5A-CEBPB-PML, evidences have been shown for the STAT5A-CEBPB interaction: STAT5A was demonstrated to cooperate with CEBPB to regulate gene transcription[64]; STAT5A can induce deacetylation of CEBPB[65]. Their interaction with PML is less well studied but STAT5 is shown to be activated by PML/RARα fusion protein in acute myeloid leukemia[66]. These common cliques in both GM12878 and K562 indicate their cell-type independent cooperation.

We next investigated whether these common cliques bind to the same loci in the two cells. For each clique, we identified the sites bound by all the member DBPs and counted how many of them are shared between the two cell types. We observed that the CTCF-RAD21-SMC3 clique shared 13,960 (51.3%) common binding sites in K562 and GM12878 (Fig. 3g), which is in agreement with the general roles of CTCF and cohesin complex in stabilizing loops[29]. The MAX-MYC-MAZ-CHD2 clique shows moderate conservation with 729 (15%) common binding sites across the two cell types. In contrast, the binding sites of P300-H3K4me1-H3K4me2 and STAT5A-CEBPB-PML are highly cell-type specific. Since P300-H3K4me1-H3K4me2 mark active enhancers and enhancers are highly cell type specific, it is understandable that there are only 62 (1.2%) of P300-H3K4me1-H3K4me2 peaks are shared across cell types. Only a small percent (2.4%, 117 sites) STAT5A-CEBPB-PML sites shared between GM12878 and K562 was unexpected. To investigate the reason why STAT5A-CEBPB-PML has distinct binding profiles in the two cell types, we first analyzed the enriched GO terms for the sites bound by all three DBPs in K562 and GM12878 respectively. Enriched GO terms in each cell type are highly specific: the top terms in GM12878 are “immune response” and “lymphocyte activation”; sites in K562 are enriched with GO terms such as “platelet activation” and “erythrocyte differentiation” which are highly specific to K562.

The above analyses showed that the same DBPs can bind to different loci to regulate cell type specific functions, which is likely due to their cooperation with different partners in different cell types. To identify their cell-specific partners, we examined the binding peaks of all the available ChIP-seq data in the regions bound by STAT5A-CEBPB-PML in GM12878 and K562 (Fig. 4a and Fig. 4b). It is obvious that the co-occurring DBPs are completely different in the two cell types: RUNX3, ATF2, MEF2A in GM12878 compared to TEAD4, TAL1, GATA2 in K562. Because of the limited number of ChIP-seq experiments, possible partners might not be profiled. Therefore, we performed de novo motif analysis using MEME-ChIP[67] in STAT5A-CEBPB-PML sites and then matched the found motifs to the known ones. It is also obvious that the de novo motifs found in K562 and GM12878 were very different. Encouragingly, all de novo motifs matched to the motifs recognized by the DBPs showing co-occurring ChIP-seq peaks. These results suggest STAT5A-CEBPB-PML complex indeed have different regulatory mechanisms in different cell types. To further interrogate the regulatory mechanisms of their cooperation, we used Spamo[68] to find spacing constraints between de novo motifs. As a result, in GM12878 we found two de novo motifs, corresponding to STAT5 and MEF2, showing a statistically significant spacing constraint with MEF2 motif occurring 13 bp downstream of STAT5 motif (Fig. 4c). This finding is novel as there is no previous report about the partnership of STAT5 and MEF2. On the other hand, in K562 the TAL1::GATA1 motif frequently appears at the upstream of RUNX1 sites at a distance of 38 bp. GATA and RUNX usually cooperate with each other and form a cis-regulatory module[69–71]. Therefore, our analysis has identified both novel and known spacing constraints between TFs.

**Figure 4.**
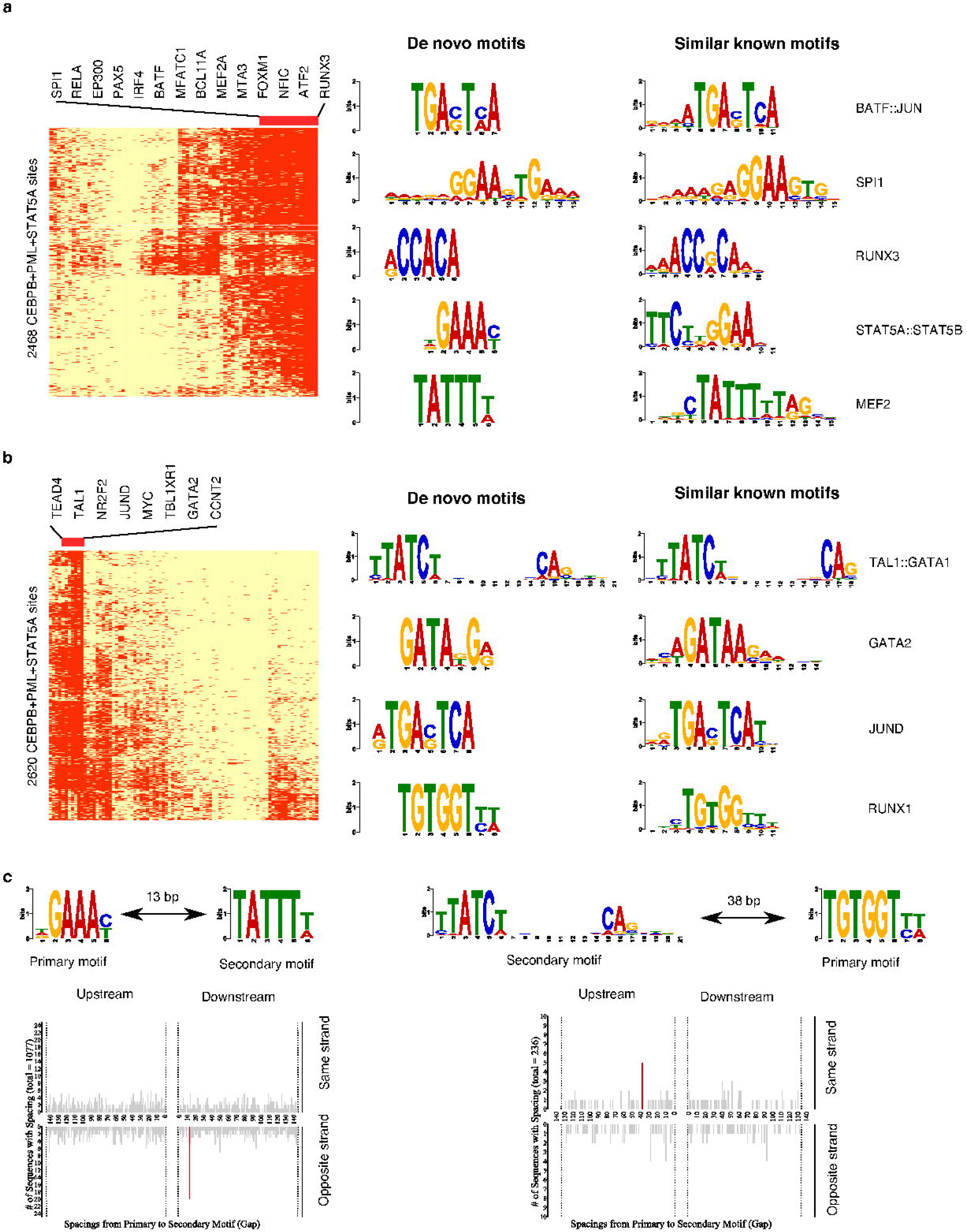
CEBPB-PML-STAT5A cooperates with different DBPs in GM12878 and K562. (a) DBP binding profile (Left) and enriched de novo motifs (Right) in 2468 CEBPB-PML-STAT5A binding sites in GM12878. (b) DBP binding profile (Left) and enriched de novo motifs (Right) in 2620 CEBPB-PML-STAT5A binding sites in K562. (c) Enriched spacing between de novo motifs found in CEBPB-PML-STAT5A sites in K562 and GM12878.

## Discussion

We present here a novel method to systematically identify both cis and trans cooperation between DBPs the first time. Many of these interactions are likely resulted from physical interactions as most of the edges in the DBP cooperation network are supported by the PPI data. Our results showed that trans-cooperation between TFs is ubiquitous, indicated by 86% of identified associations having strong trans correlation scores, which can only be discovered by integrating DBP binding and chromatin structure data. These trans-interactions most often accompany cis-interactions as the majority (71%) of DBP cooperation is a mixture of trans and cis events. Furthermore, we observed enrichment of disease associated GVs in DBP cooperative binding sites, which suggests the functional importance of DBP interactions.

Identification of cooperation between multiple DBPs has been a challenging problem. Combinatorial approaches are limited to consider cooperation between small number of DBPs because of the exponential increase of the possible combinations. In contrast, our model can easily search combinatorial cooperation in thousands of DBPs. By identifying modules and cliques in the network, we have uncovered closely collaborated DBPs, particularly those associated through trans interactions in chromatin loops that may be crucial for loop formation or stabilization.

Our comparative analyses between GM12878 and K562 revealed different mechanisms of achieving cell-type specificity: using different combinations of DBPs or using the same protein complex but collaborating with different partners. Interestingly, we also found spacing constraints between the binding sites of certain partners, which implies higher-order of regulatory rules for not only cis but also trans DBP cooperation. As ChIP-seq data and HiC data are rapidly accumulating, our model provides a powerful tool for integrative analysis of DBP binding and chromatin structure data in different cell types, which will facilitate to uncovering the molecular mechanisms for transcriptional regulation and 3D chromosome organization.

## Methods

### Data sets

BAM files of ChIP-seq experiments in K562 and GM12878 were downloaded from the ENCODE project website[72]. Chromatin loops were downloaded from the Rao et al. study[29]. Because only these two cell lines had both DBP ChIP-seq and 5Kbp-resolution Hi-C data, we focused on these data sets in this study.

### Data Preprocessing

We divided each chromosome into consecutive 1 kb regions. For each protein, we computed the Reads Per Kilobase per Million mapped reads (RPKM)[24] on these regions. The fold enrichment was calculated using MACS’s algorithm[73] with customized parameters. In particular, we set λ_local_ = max(λ_BG_, λ_14k_, λ_24k_, where λ_BG_ is the average RPKM of the whole genome; λ_14k_ and λ_24k_ are average RPKM of 14 kb and 24 kb windows. We used a larger window size than MACS’s default size and a loose p-value (0.01) to call peaks. By doing so, we increased the sensitivity for detecting broad peaks. For each 1 kb region, if it was called as a peak, we used the fold enrichment as its ChIP-seq enrichment score; otherwise, a zero score is assigned to that region. After computing the enrichment scores for every protein, we removed regions with low variation of scores by requiring the standard deviation of scores is at least 1. This excludes some unwanted artifacts from our analysis. For instance, regions with low mappability or abnormal high signal[72]. Next, for each DBP pair we calculated the Spearman’s correlation of ChIP-seq enrichment scores in the remaining bins as the cis-correlation score. To compute trans correlation scores, we first downloaded the 3D interaction loops identified in a 5Kbp-resolution Hi-C study[29]. Next, for each DBP we computed its enrichment scores on loop regions as follows: Suppose we have n loops, denoted by L^(1)^, L^(2)^,…, L^(n)^ and each loop L^(i)^ consists of two interacting loci 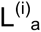 and 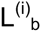. To compute the enrichment score of a given DBP on loop L^(i)^, we first binned 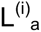 into 1 kb regions, and then take the maximum of ChIP-seq enrichment scores of these bins as the enrichment score for 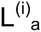. Likewise, we can compute the enrichment score for 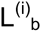. Then, given a pair of DBPs denoted by A and B, for every loop, we first compared the enrichment scores of A on the two interacting loci, and took the larger one as the primary binding locus of A. As oppose, we assume the smaller one to be the primary binding locus of B. The enrichment scores of primary binding loci for each protein were then used to compute correlation coefficient.

### Network Construction

We adapted the Gaussian graphical model (GGM) to construct the DBPs cooperation networks. GGM assumes that the observations have a multivariate Gaussian distribution with mean μ and covariance matrix Σ. Let Σ^−1^ be the inverse of covariance matrix. If the ij th component of Σ^−1^ is zero, then variable i and variable j are conditional independent given other variables. Therefore, each non-zero component represents an edge in the network. To efficiently and accurately estimate the inverse of covariance matrix using DBP ChIP-seq data, we employed the Graphical lasso algorithm[20] and the Copula method [21, 22]. Specifically, we used the subroutine “huge” from the “huge” R package[74], with the lasso penalty equal to 0.3. We choose this value since less than 15% of DBP pairs have correlation score greater than 0.3. In addition, most of the known interactions are above 0.3. Therefore, we decided 0.3 is an appropriate value from our experience. Because we aimed at identifying DBP interactions, edges with negative correlations were removed in the network analysis.

### Network analysis

We used Eppstein’s algorithm[75] for maximal clique searching, which gives an exact solution in near optimal time. For community detection, we used Newman’s algorithm[47]. Both implementations were provided in the “igraph” C library[76].

### Comparing DBP cooperation network with protein-protein interaction (PPI) network

Protein-protein interaction data was obtained from the BioGrid database[77] version 3.2.99. For each edge formed by node A and B in a DBP cooperation network, if this edge is also present in the PPI network, it was considered as a direct interaction. Otherwise, we checked whether there exists a third node in the PPI network that connects to both A and B; if so, this edge was considered as an indirect interaction. To determine the statistical significance of these overlaps, we first replaced the nodes in the DBP cooperation network with randomly selected genes from the PPI network. Next, we counted the direct and indirect interactions in the simulated network. This process was repeated for 10^9 times to generate the background distribution, which was then used to calculate the p-values.

### Simulated networks

To generate a Erdős–Rényi random graph, we used the G(n,p) model[78]. This model specifies an n-node network, in which each edge is included with a probability p independent from every other edge. We used p = 0.2 in this paper which gives rise to a sparse network. We follow the procedures in ref [22] to generate a Gaussian distributed dataset which was used for constructing the simulated network.

We used Genetweaver 3.1 to extract random subnetworks with different sizes (50 and100 nodes) from the yeast gene regulatory network provided by the software. For other parameters, we used the software’s default setting.

To draw the ROC curve, we counted the number of true positives (TPs), false positives (FPs), true negatives (TNs) and false negatives (FNs). If a predicted edge is present in the true network, it is a TP, otherwise it is a FP. For edges that are present in true network but not identified by the algorithm, they are FNs. TNs are edges that are not present in either predicted or true networks.

## Supplemental data

Supplemental Table 1: Speed comparison of GGM and ARACNE on data sets with different sizes.

Supplemental Table 2: Centrality of nodes in GM12878 DBP cooperation network.

Supplemental Table 3: A list of cis, trans and mix associations in GM12878.

Supplemental Table 4: Fours communities found in the GM12878 DBP cooperation network.

Supplemental Table 5: All DBP cliques found in GM12878.

Supplemental Table 6: Cell type specific DBP cliques and common DBP cliques in GM12878 and K562.

Supplemental Figure 1: Degree distribution of DBP cooperation network in GM12878.

Supplemental Figure 2: Functional analysis of communities in the GM12878 DBP cooperation network. (a) Enriched GO terms in green community. (b) Enriched GO terms in cyan community. (c) Enriched GO terms in red community.

